# CRAM compression: practical across-technologies considerations for large-scale sequencing projects

**DOI:** 10.1101/2022.12.21.521516

**Authors:** Ayesha Al Ali, Palani Kannan Kandavel, Hakeem Al Mabrazi, George Carvalho, Vinay Kusuma, Gurunath Katagi, Santosh Elavalli, Ayman Yousif, Mohammad Riyaz Akhter, Joseph Mafofo, Tiago Magalhaes, Javier Quilez

## Abstract

CRAM is an efficient format to store high-throughput sequencing data and it has been widely adopted. We thus plan to use CRAM for the Emirati Genome Program, which aims to sequence the genomes of ~1 million nationals in the United Arab Emirates using short- and long-read sequencing technologies (Illumina, MGI and Oxford Nanopore Sequencing). We conducted a pilot study on the three technologies before start using CRAM at scale. We found CRAM achieved 40–70% compression depending on the sequencing platform. As expected, CRAM compression was data lossless and did not alter variant calls. In our cloud, we observed compression speeds 0.7–1.4 GB per minute, varying on the sequencing platform too. This translates into ~1–2 hours using a single CPU to compress a ~30X human whole-genome sequencing sample. Despite its wide use, we found little publicly available information about CRAM compression rate, speed, losslessness and parallelization, especially across many sequencing platforms. This work will have direct application for Emirati Genome Program and provide practical considerations for other large-scale sequencing efforts.

## Introduction

High-throughput sequencing (HTS) and the associated bioinformatics analysis results in several large files (Supplementary Note 1 and Supplementary Table 1). As an example, storing all key files generated for a human ~30X whole-genome sequencing (WGS) sample (i.e. raw format, FASTQ, BAM and gVCF) amounts to ~200–900 gigabytes (GB) depending on the sequencing technology – this amounts to ~50–250 USD of storage per year in AWS. This poses an infrastructure and cost challenge for large-scale genome sequencing efforts such as the Emirati Genome Program (EGP). In EGP, we will sequence at ~30X all 1 million nationals in the United Arab Emirates (UAE). EGP started in 2021 and it is expected to be completed by 2025. As of November 2022, we have sequenced the genomes of >250,000 EGP participants at ~30X employing three HTS platforms (Illumina, MGI and Oxford Nanopore Technologies; the latter hereafter referred to as “ONT”). To quantify the infrastructure and cost challenge, we estimate that a maximum of ~140 petabytes (PB) of storage would have been required to store raw and processed data for all ~250,000 genomes if those *co-existed* at the same time.

Therefore, a well-thought strategy on which files are stored in the long term to minimize the storage footprint while meeting the project requirements is crucial. This is not only especially important for sequencing projects with high volumes of samples. Besides, it is also for those in which re-analysis of the data is expected and / or with contractual obligations to be able to deliver not only the genetic variants but also some form of raw data (e.g. sequencing reads). Several well-known worldwide organizations and sequencing projects have chosen a long-term storage of sequencing data in the form of CRAM files. Given that, for EGP we advocated a strategy consisting in the long-term storage of only CRAM and (g)VCF files (see Supplementary Note 1 for more details) upon successful completion of the pipeline and confirmation that the target quality control (QC) metrics are met.

Before implementing such strategy, we conducted a proof-of-concept (POC) study using 30 genomes sequenced on the three sequencing platforms in our G42 Healthcare’s Omics Center of Excellence (Illumina, MGI and ONT; 10 genomes per platform). In the POC, we assessed the feasibility, compression rate, data lossless, speed of the BAM-to-CRAM compression and optimal parallelization across the three platforms. Here we present the results of that POC and expand with practical considerations for conducting BAM-to-CRAM compression at scale as well as to directly generate CRAM files with existing analysis pipelines. Besides, we discuss on the projected cost savings resulting from using CRAM, its limitations and existing alternatives. To our knowledge, an exercise like this has not been published or made available to the community. Therefore, we expect this work to be useful for sequencing projects concerned about effectively minimizing the storage footprint with little or no impact on their analysis pipelines.

## Results and Discussion

Our dataset consisted of 30 distinct human genomes sequenced on two short-read (Illumina and MGI) and one long-read (ONT) sequencing technologies (10 genomes on each platform). We sequenced and analyzed the 30 genomes as described in the **Materials and Methods**, resulting in 30 BAM files with per-genome average coverage between 20 and 120X (Supplementary Table 2).

### CRAM achieves 40–70% compression depending on sequencing platform

We first aimed to replicate in our sequencing and analysis setup the compression rates observed by previous authors as well as to determine if those generalize across sequencing platforms. Specifically, we converted each of the 30 BAM files into CRAM format using Samtools (Li et al., 2009) and, for each file, we calculated the compression rate as the size of the file size reduction relative to the original BAM (see **Materials and Methods**). We observed compression rates between 40% to 70%, varying depending on the sequencing platform (Figure 1a). Such values are in the same range as previously reported for Illumina data (Bonfield, 2022). Compression rates for short-read data (65.7% and 50.7% for Illumina and MGI, respectively) were higher compared to that of long-read ONT (39.6%). We hypothesize that the constant read length nature of short-read data makes compression easier compared to the variable read length of ONT. We think multiple factors can explain the differences in compression rate between the two short-read fixed-length platforms. We initially speculated that the longer read length used in this POC for Illumina (150 bp) compared to MGI (100 bp) increases the compression rate of the former. However, we observed similar compression rates (~50%) in 6 MGI samples sequenced each with both 100 and 150 bp flow-cells (data not shown). Illumina versus MGI differences might be also partly explained by the longer FASTQ headers we observed in MGI compared to Illumina (78 and 66–68 characters, respectively). Besides, the hard trimming of reads we applied to MGI reads to remove sequencing adapters and low-quality ends leads to some minimal variation in read length distribution. Such variation would not be present in soft-clipped reads processed with DRAGEN, potentially contributing to a higher compression rate. Finally, we observed higher duplication rates for Illumina data compared to PCR-free MGI data (data not shown), which may facilitate compression of the former and hence its higher compression rate. We also observed that per-BAM compression rate is independent of the input BAM size (Figure 1b), which simplifies estimating the file size reduction that can be achieved with CRAM compression. Of note, we found quite remarkable that FASTQ and BAM files generated from MGI data are both about two times larger than those generated from Illumina and even ONT (Supplementary Table 1 and Supplementary Table 2). We wonder whether this is again due to read length differences, similarly as seen for compression rate of Illumina versus MGI samples (Figure 1a), or intrinsic to the MGI sequencing technology. If MGI intrinsically yields to larger FASTQ and BAM files as well as lower CRAM compression rates, it is worth considering the lower storage footprint of Illumina compared to MGI.

**Figure 1.**
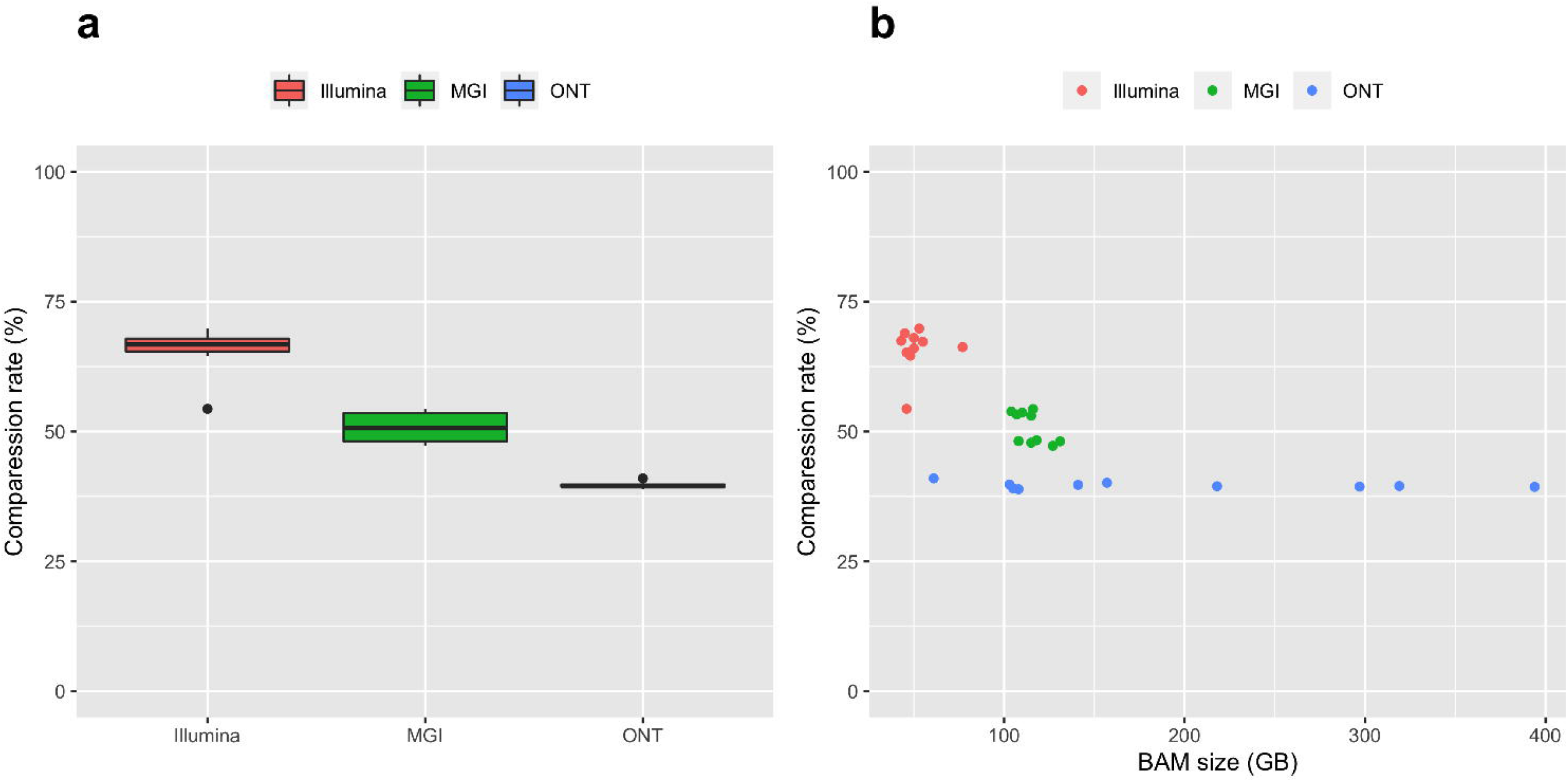
Compression rate across the three platforms. **(a)** Compression rate distributions across platforms, with average values of 65.7% (Illumina), 50.7% (MGI) and 39.6% (ONT). **(b)** No relationship between original BAM size (GB) and compression rate.

### CRAM compression is data lossless and does not alter variant calls

Because for each genome we had FASTQ, BAM and VCF files, we could confirm that converting from BAM to CRAM did not alter sequencing data at different processing stages. Firstly, for each CRAM we regenerated FASTQ files (one for each read of the pair) and determined three key QC metrics on these: total number of sequenced gigabases (Gb), average sequencing read and average Q30 score. When compared to the same QC metrics generated on the starting FASTQ files (i.e. prior to BAM generation through alignment), we observed 100% similarity in all 30 genomes. Besides, we used the generated CRAM files as input for variant calling. For each genome, the resulting VCF showed the same number of variants (even when broken into different types of variants, e.g. SNP, INDEL, etc.) and 100% concordance compared to the VCF obtained from the original pipeline. Through this experiment we noted an important consideration when re-generating FASTQ files from BAM or CRAM in paired-end sequencing samples. The sequencing reads in each of the two resulting FASTQ files (one for each read of the pair) will be by default sorted based on the genomic coordinates in the BAM or CRAM. This virtually always will result in read1 and read2 not being in the same order for exactly all pairs. As many aligners expect proper matching of read1 and read2, sorting by read name (e.g. with Samtools) before using the regenerated FASTQ files is needed.

### Optimization of BAM-to-CRAM conversion speed

Finally, we aimed to determine the speed at which we could convert BAM to CRAM format while optimizing parallelization and efficiency. As baseline, we measured BAM-to-CRAM conversion speed for each sample, defined as the number of GB compressed per minute using a single CPU (Linux machine with 64 vCPUs and 256 GB). We observed a reverse trend across platforms (Figure 2) compared to the compression rate values (Figure 1). Our explanation is that the higher the compression rate that is achieved, the more time is required to achieve it and therefore the lower compression speed. Considering the observed platform-specific BAM sizes (~75–125 GB) and compression speeds (0.7–1.4 GB / min), we estimate that compressing the BAM file from a ~30X human genome takes approximately 1–2 hours using a single CPU. We then repeated the BAM-to-CRAM compression with increasing number of CPUs and calculated the acceleration relative to a single CPU as well the efficiency (see **Materials and Methods**). As shown in Table 1, we observed a 4X increase in compression speed when using 4 CPUs instead of one, as expected for a perfect parallelization efficiency. However, as more CPU per samples are used, efficiency decreases. Of note, allocating 32-times more resources only results in completing the compression 9-times faster. Altogether, we concluded that using 4 CPU per BAM file is optimal. Our estimates of the platform-specific compression speed and optimal CPU allocation are useful to project time to completion of high volumes of BAM files to be compressed.

**Figure 2.**
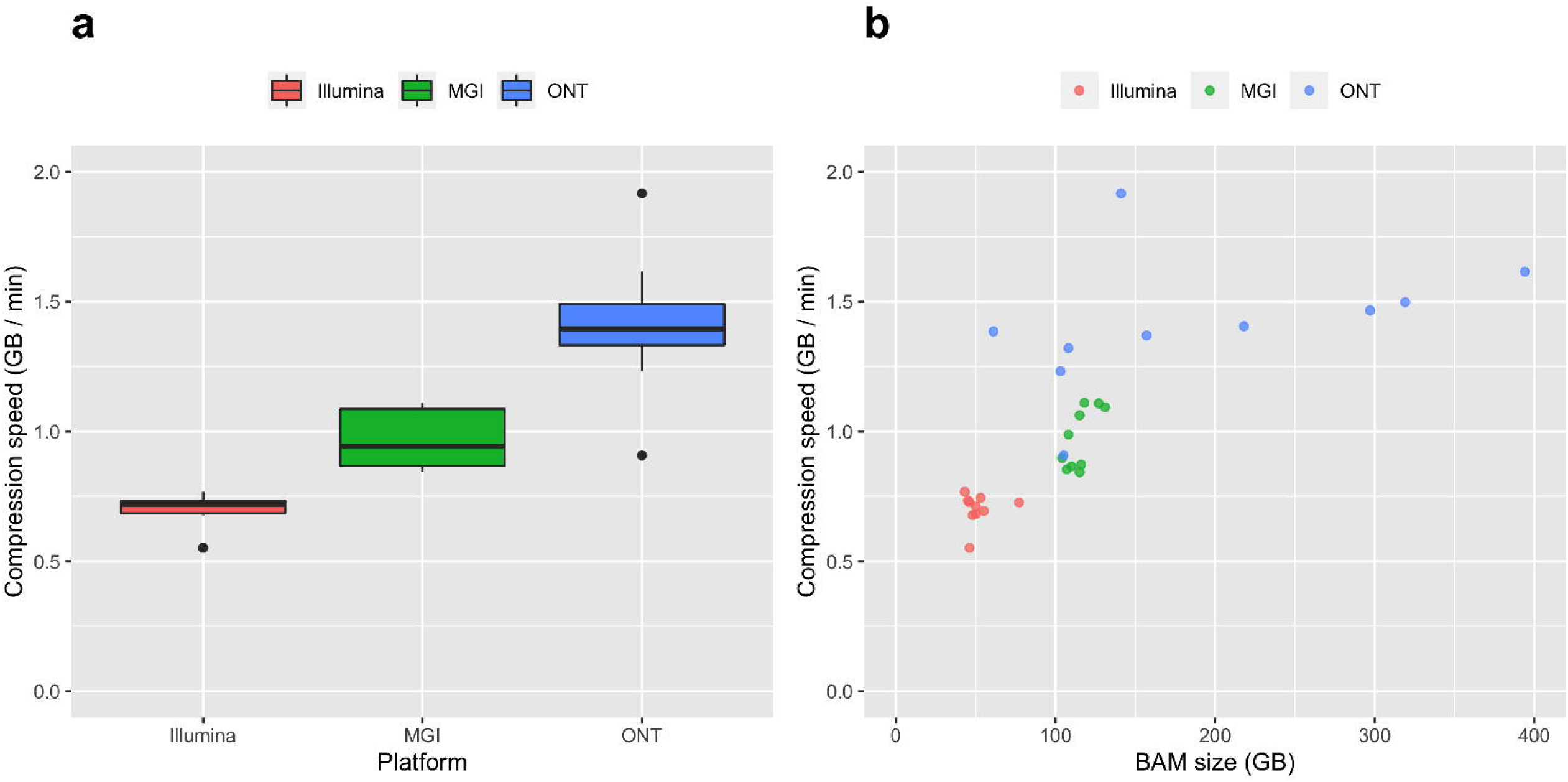
Compression speed across the three platforms. (a) Compression speed distributions across platforms, with average values of 0.70 (Illumina), 0.97 (MGI) and 1.41 (ONT) GB per minute. **(b)** Relationship between original BAM size (GB) and compression speed.

**Table 1.** Estimating optimal CPU usage for BAM-to-CRAM compression. Average speed, acceleration and efficiency for different CPU threads based on 6 samples (2 from each sequencing platform).

| Parameter | 4 CPUs | 8 CPUs | 16 CPUs | 32 CPUs |
| --- | --- | --- | --- | --- |
| BAM-to-CRAM compression speed (GB / min) | 3.85 | 7.14 | 9.09 | 9.09 |
| Acceleration relative to 1 CPU | 4.05 | 7.30 | 9.00 | 9.06 |
| Efficiency | 1.01 | 0.91 | 0.56 | 0.28 |

### Beyond this POC

Converting existing BAM files generated for EGP into CRAM will importantly reduce storage stress to our cloud and save costs. As of November 2022, we estimate that EGP-generated BAM files alone occupy ~10 PB of our cloud storage. This is calculated from the split of EGP genomes across the three sequencing platforms and a conservative (+25% buffer) BAM file size we observe per platform for a ~30X WGS (Illumina and ONT: 75 GB; MGI: 150 GB) (Supplementary Table 1). We estimate the resulting CRAM will use instead ~4 PB in our cloud (~58% storage footprint reduction), representing an annualized cost saving of ~1.2 million USD. We came up with some practical considerations for implementing CRAM compression for thousands of BAM files. In the POC, we assessed CRAM data integrity by comparing both (i) key metrics in the original FASTQ and that re-generated from the CRAM and (ii) variant calls generated from BAM and CRAM. Such sanity checks are time consuming and hence impractical at scale. Instead, we argued that a data lossless CRAM will have a Samtools flagstat file identical to that of the source BAM. Therefore, we suggest requiring for identical Samtools flagstat between the original BAM and the generated CRAM on the fly is an efficient and fast strategy to confirm CRAM compression was data lossless. Going forward, we plan to directly generate CRAM files instead of compression from BAM files. The DRAGEN (Illumina, 2021) and Sentieon software we use for Illumina and MGI data, respectively, can directly generate and read CRAM files. Clair3 (Zheng et al., 2022), the variant caller we use for ONT data, cannot read CRAM yet so we plan to convert BAM to CRAM once variant calling completes. In any case, CRAM has some limitations and commercial solutions like PetaGene can achieve similar compression rates and easier functionality at a cost (Supplementary Note 1).

## Materials and Methods

### HTS data processing pipelines

#### Illumina

The BCL file is the native output format of Illumina sequencing systems. We used on-premise Illumina DRAGEN Bio-IT Platform 3.9 (Illumina, 2021) to de-multiplex and base-call BCL files into per-sample FASTQ files. We also used Illumina DRAGEN Bio-IT Platform 3.9 to align reads in FASTQ files to the GrCH38 human reference genome, post-alignment processing (sorting and marking duplicates; base quality score recalibration), call genetic variants and generate summary statistics.

#### MGI

The CAL files is the native output of MGI sequencing systems. We used on-premise MGI Ztron server and Zebra software to de-multiplex and base-call CAL files into per-sample FASTQ files. Such files were pushed to G42 Cloud for initial QC assessment as well as trimming of sequencing adapters and low-quality ends with fastp (v0.23.2) (Chen et al., 2018). We aligned the trimmed FASTQ files to the GRCh38 human reference genome using the BWA-MEM algorithm (Li and Durbin, 2010) implemented in Sentieon (sentieon-20211202). We used alignments in BAM format for marking duplicates, estimating effective coverage and call SNP and INDEL variants using Sentieon as well (haplotype caller algorithm). We converted the resulting gVCF into VCF an calculated key metrics for the SNP and INDEL calls.

#### ONT

P48 sequencing results in many Fast5 files for each sample, which we processed in G42 Cloud using an in-house custom pipeline including ONT-recommended tools. Firstly, we base-called the Fast5 files with ONT’s proprietary tool “Guppy” [v4.4.1, v.6.1.3]. This resulted in as many FASTQ files as Fast5 files used as input, which we merged to have a single FASTQ file per sample. On each sample we ran MinIONQC (Lanfear et al., 2019) to perform the initial sequencing QC and check if the target total number of Gb was achieved during the sequencing. We aligned FASTQ files to the GRCh38 human reference genome using Minimap2 (Li, 2018) and we used Alfred (Rausch et al., 2019) to check the quality and alignment QC for each EGP sample. The alignments generated by Minimap2 were stored in BAM format, which we later converted into CRAM. We used the alignments (in CRAM format after converting from the BAM generated by Minimap2) to call SNP and INDELs as well as structural variants (SV) using Clair3 (Zheng et al., 2022) and Sniffles2 (Sedlazeck et al., 2018), respectively. We used VariantQC (Yan et al., 2019) for performing the quality checks on the variants called and reporting the statistics for each sample.

### CRAM compression

#### Samtools and CRAM 3.1 specification

We used CRAM v3.1, an improvement of CRAM v3.0, which provides more reduction in file size. Specifically, Illumina CRAM 3.1 is 7% to 15% smaller than the equivalent CRAM 3.0 and 50% to 70% smaller than the original BAM file (Bonfield, 2022). We used Samtools (Li et al., 2009) v1.15 available in GitHub for CRAM compression from BAM as well as for CRAM file manipulations. We chose the latest version of such tool when we conducted the POC because it supported CRAM v3.1.

#### Compression rate

We defined compression rate as the ratio of size reduction relative to BAM size (or in general, uncompressed data) (see Equation 1).

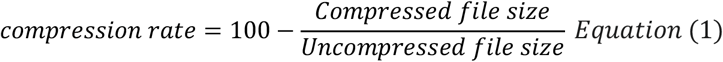

#### Data integrity and losslessness

The CRAM compression is lossless and allows restoring the original data without any loss of information. Firstly, we assessed whether sequencing reads remained unaltered after CRAM compression from BAM. By uncompressing CRAM files to FASTQ, we ran fastp (Chen et al., 2018) on the latter to determine key QC metrics for each sample: (i) total number sequenced Gb, (ii) average sequencing read length, and (iii) average Q30 score. The three quality parameters are set to ensure lossless compression and validate intact data of each sample. As a result, for all samples we had no loss in all the parameters. Secondly, we wanted to confirm that CRAM compression did not change the genetic variant calls. Specifically, we performed variant calling as described elsewhere in the manuscript using either the original BAM or the post-compression CRAM file as input. We compared (i) the total number of SNPs, indels, multiallelic sites, and multiallelic SNP sites as well as (ii) the genomic position and genotype concordance. We did the latter by pairwise comparison of the VCF files with BCFtools (Danecek et al., 2021) and calculating the concordance as the Jaccard index (Equation 2).

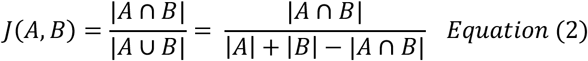

#### Tools and command lines

BAM to CRAM command

~~~
samtools view -C -T ${referenceFile.fna} ${bamFile} -o ${outDir}/${sampleID}.cram -@8
~~~

Data integrity check

~~~
bamFlagstat=‘md5sum <(samtools flagstat ${bamFile} -@ 8) | cut -f1 -d“ ”’
cramFlagstat=‘md5sum <(samtools flagstat ${cramFile} -@ 8) | cut -f1 -d“ ”’
if [[ “$bam” == “$cram” ]]
then
  echo “matched flagstat for ${sampleID}”
else
  echo “unmatched flagstat for ${sampleID}”
~~~

CRAM to FASTQ

~~~
samtools fastq --reference ${referenceFile.fna} -1 file1.R1.fastq -2 file2.R2.fastq
${cramFile.cram}
fastp -A -G -Q -L -w 1 -i file1.R1.fastq -I file2.R2.fastq -h output_fastp.html
~~~

VCF files comparison

~~~
bcftools isec <A.vcf.gz> <B.vcf.gz> -p <dir>
~~~

#### Optimizing resources

Optimal speed (Equation 3) of compressing BAM to CRAM is achieved by parallelizing the compression process and determining the optimal computing resources utilization. To determine the optimal number of CPUs per sample, we calculated the acceleration of multiple threads relative to one thread using Equation 4 and the efficiency of utilization using Equation 5. We generated the results in the previous sections by using 1 CPU to compress each sample. For 2 out of the 10 samples from each platform, we repeated CRAM compression with increasing CPU threads 4, 8, 16, and 32 threads.

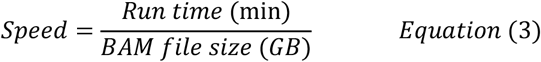

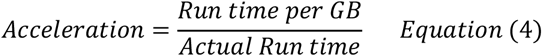

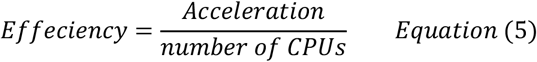

## Acknowledgements

NA.

## Authors Contributions

Study design: AA, JQ, PKK. Sequencing: MRA, JM. Infrastructure and processing of sequencing data: AA, HAM, GC, VK, GK, SE, AY, PKK. Data analysis: AA, JQ. Manuscript writing: AA, JQ, TM.

## Supplementary Information

**Supplementary Table 1.** File types and sizes generated across Illumina, MGI and ONT sequencing platforms. The table shows file sizes for ~30X WGS samples as an average across multiple EGP samples. For Illumina and MGI those corresponded to 150- and 100-bp paired-end read lengths, respectively. To be conservative, the FASTQ and BAM file size estimates in the table are increased by 25%. Likewise, we rounded up VCF and gVCF to 1 and 10 GB, respectively. CRAM file size estimates are calculated by multiplying the BAM file size in the table by the average compression rate obtained for each sequencing platform in this POC.

| Sequencing platform | Raw signal | Sequencing reads | Alignments | Variant calls | Variant calls |
| --- | --- | --- | --- | --- | --- |
| Illumina | BCL<br>(50 GB<sup>1</sup>) | FASTQ<br>(75 GB) | BAM<br>(75 GB) | gVCF<br>(10 GB) | VCF<br>(1 GB) |
| MGI | CAL<br>(150 GB<sup>1</sup>) | FASTQ<br>(150 GB) | BAM<br>(150 GB) | gVCF<br>(10 GB) | VCF<br>(1 GB) |
| ONT | Fast5<br>(700 GB) | FASTQ<br>(75 GB) | BAM<br>(75 GB) | gVCF<br>(10 GB) | VCF<br>(1 GB) |

<sup>1</sup>Calculated as BCL file size divided by the number of multiplexed samples.

### Supplementary Table 2. Sequencing throughput and BAM files sizes for the 30 genomes included in the POC

“Supplementary Table 2.xlsx”

### Supplementary Note 1. HTS file types and long-term storage strategy

HTS raw and processed data consist of standard relatively large files (in the order of GB). Size-reduction common practices exist for most of them and, yet, storing all file types is redundant and impractical. Here we briefly describe such file types and size-reduction practices as well as we discuss on long-term file type storage.

#### HTS raw and processed file formats in Illumina, MGI and ONT

Most sequencing technologies have a proprietary format to store the raw data generated by their sequencing instruments. For instance, sequencing-by-synthesis technologies such Illumina and MGI use image-based BCL and CAL files, respectively, to store data generated by their instruments. ONT relies on the HDF5 format as the base for the Fast5 files storing the electrical signal generated by their long-read sequencers. Through base-calling (commonly referred to as “primary analysis”), such technology-specific file formats converge into the standard FASTQ format to store the read sequence and quality scores. Likewise, the “secondary analysis”, which for many sequencing applications comprise alignment to a reference sequence and variant calling, employ standard formats across platforms. For instance, alignment software tools express alignments following the SAM format specifications (Li et al., 2009). Finally, variant callers write genetic variants in VCF format (Danecek et al., 2011) (or the related gVCF format when merging across multiple genomes is planned or for certain downstream tools). Due to the relatively large size of the files above, common practices exist to reduce the storage footprint of such files. For instance, FASTQ files are preferably gzip-compressed, which, as a rule-of-thumb, reduces file size by 50%. Besides, the .ORA format developed by Illumina is 5-times smaller relative to the original FASTQ. Alignments are rarely stored in SAM format but written and read in its binary BAM format instead; furthermore, BAM files can be compressed into CRAM format for an additional ~50% file size reduction relative to the former. Finally, VCF files are typically “gzipped” when generated by variant callers and can be in its binary format (“.bcf”) for additional file size reduction, with both formats being accepted by most bioinformatics tools. Despite the approaches above to reduce the size of each file type, storing all file types generated in the analysis pipeline is redundant and, more importantly, may be impractical due to the tremendous storage footprint (especially for large-scale sequencing projects and / or those required to store data for a long time).

#### Thoughts on efficient long-term storage strategies

##### Platform-specific raw data (BCL, CAL and Fast5)

Raw data like BCL or CAL files are very rarely kept upon certain relatively short time or when key QC metrics are passed, especially considering the relatively big size of such files and the fact that the base-calling process is mature and little-changing for Illumina and MGI sequencing. Conversely, more difficult is the decision of deleting ONT’s Fast5 files. For one, ONT’s sequencing technology as well as the base-callers software tools and used deep learning models are more frequently evolving to catch up with the lower error rates of other technologies like Illumina, MGI and PacBio. Such ongoing improvements not only occur for the “canonical” base-calling (i.e. determination of DNA sequence) but for its capability to infer multiple methylation changes (e.g. 5mC, 5hmC) from the electric signal in the Fast5 files, a key competitive edge of ONT relative to those competitor technologies. Altogether, deleting the heavy Fast5 files (~700 GB for a 30X human genome) is unavoidably accompanied by the fear of higher-accuracy and data for additional methylation marks if kept.

##### Raw sequencing reads (FASTQ and .ORA)

Some argue to keep FASTQ files as long-term storage of sequencing data, some of the reasons being that (i) it is closer to the raw data (e.g. no trimming of sequencing adapters and / or low-quality read ends), (ii) re-alignment may be needed with improved aligners, and (ii) FASTQ files are the starting point for different applications, i.e. not only alignment plus variant calling. In this regard, Illumina advocates for storing sequencing reads in its .ORA format is smaller even compared to the CRAM format. This has some disadvantages: (i) .ORA does not contain alignment information so the resource-consuming alignment step will need to be repeated in most re-analyses; (ii) Illumina’s proprietary DRAGEN software is required to read .ORA files, potentially creating vendor lock-in; (iii) analysis workflow may be complicated if .ORA files are used for long-term storage in the cloud and any re-analysis needs to be done with on-premise DRAGEN units.

##### Alignments (BAM and CRAM)

We see key advantages in using the alignments (BAM, CRAM) compared to the raw sequencing reads (FASTQ, .ORA): (i) additional information derived from the alignment is available or can be calculated (e.g. coverage); (ii) resource-consuming alignment process is not needed in the event some re-analysis on the BAM is required; (iii) most aligners can use BAM format as input for re-alignment; and (iv) even for those which do not, BAM can be converted to FASTQ or “piped” to any tool requiring FASTQ as input with tools such as Samtools (Li et al., 2009). CRAM has limitations too. For instance, the reference FASTA used during the compression is required to re-generate the BAM from the CRAM, which is moreover a time-consuming task. Besides, not all tools which accept BAM can accept CRAM too and we think that BAM is still the default preference for alignments by many, relatively limiting the spread of CRAM. PetaGene offers a commercial solution that overcomes some of such limitations.

##### gVCF and VCF files

(g)VCF files are unlikely to be deleted because are the end point of the primary and secondary analysis, are used for downstream analyses and are relatively light anyway.

